# Exploiting a metabolic vulnerability in brain tumour stem cells using a brain-penetrant drug with safe profile

**DOI:** 10.1101/2024.01.15.574967

**Authors:** Audrey Burban, Cloe Tessier, Mathis Pinglaut, Joris Guyon, Johanna Galvis, Benjamin Dartigues, Maxime Toujas, Mathieu Larroquette, H Artee Luchman, Samuel Weiss, Nathalie Nicot, Barbara Klink, Macha Nikolski, Lucie Brisson, Thomas Mathivet, Andreas Bikfalvi, Thomas Daubon, Ahmad Sharanek

**Affiliations:** University of Bordeaux, CNRS, IBGC, UMR5095, Bordeaux, France; University of Bordeaux, INSERM, UMR1312, BRIC, BoRdeaux Institute of onCology, Bordeaux, France; CHU of Bordeaux, Service de Pharmacologie Médicale, Bordeaux, France; University of Bordeaux, INSERM, BPH, U1219, Bordeaux, France; Bordeaux Bioinformatic Center CBiB, University of Bordeaux, Bordeaux, France; Department of Cell Biology and Anatomy, University of Calgary, Calgary, Alberta, Canada; Arnie Charbonneau Cancer Institute and Hotchkiss Brain Institute, University of Calgary, Calgary, Alberta, Canada; LuxGen Genome Center, Luxembourg Institute of Health, Laboratoire national de santé, Dudelange, Luxembourg; National Center of Genetics (NCG), Laboratoire National de Santé (LNS), Dudelange, Luxembourg; Department of Cancer Research (DoCR), Luxembourg Institute of Health (LIH), Luxembourg 1526, Luxembourg

## Abstract

Glioblastoma (GB) remains one of the most treatment refractory and fatal tumour in humans. GB contains a population of self-renewing stem cells, the brain tumour stem cells (BTSC) that are highly resistant to therapy and are at the origin of tumour relapse. Here, we report, for the first time, that mubritinib potently impairs stemness and growth of patient-derived BTSCs harboring different oncogenic mutations. Mechanistically, by employing bioenergetic assays and rescue experiments, we provide compelling evidence that mubritinib acts on complex I of the electron transport chain to impair BTSC stemness pathways, self-renewal and proliferation. Global gene expression profiling revealed that mubritinib alters the proliferative, neural-progenitor-like, and the cell-cycling state signatures. We employed *in vivo* pharmacokinetic assays to establish that mubritinib crosses the blood-brain barrier. Using preclinical models of patient-derived and syngeneic murine orthotopic xenografts, we demonstrated that mubritinib delays GB tumourigenesis, and expands lifespan of animals. Interestingly, its combination with radiotherapy offers survival advantage to animals. Strikingly, thorough toxicological and behavioral studies in mice revealed that mubritinib does not induce any damage to normal cells and has a well-tolerated and safe profile. Our work warrants further exploration of this drug in in-human clinical trials for better management of GB tumours.

## Introduction

Glioblastoma (GB) is the most common and aggressive primary tumour in the adult brain, leading to the majority of ∼200,000 deaths related to the central nervous system tumours worldwide each year^1, 2^. The current standard of care for GB patients includes surgical excision of the tumour followed by ionizing radiation (IR) and chemotherapy^3^. Despite intense efforts at targeting various signaling pathways, putative driver mutations and angiogenesis mechanisms, survival has not substantially improved beyond 18 months following diagnosis^4,5^. The main challenges underlying therapeutic failure are rooted to its inter- and intra-tumoural heterogeneity, resistance to therapy and relapse.

Intra-tumoural heterogeneity is thought to be promoted by the brain tumour stem cells (BTSCs), a population of self-renewing, multipotent, and tumour-initiating stem cells, that undergo dynamic state transitions^6, 7^. BTSCs invade extensively throughout the brain, develop a hyper-aggressive phenotype, resist therapies, and lead to tumour recurrence^8^. Therefore, developing novel approaches to suppress BTSCs are at the forefront of efforts to fight GB.

Metabolic dysregulation is one of the most important hallmarks in GB and contributes to therapy resistance^9–11^. For decades, tumour metabolism was believed to rely heavily on aerobic glycolysis, a phenomenon known as the Warburg effect^12^. However, in line with the complex heterogeneity of GB, recent reports evidenced the existence of multiple metabolic dependencies in GB where mitochondrial respiration is highlighted as an essential alternative source of energy and is shown to play a crucial role in GB tumourigenesis^13, 14^. Recent investigations have documented that GB cells also acquire functional mitochondria via tunnelling from other cancerous counterparts^15, 16^ or non-malignant host cells in the tumour microenvironment^17, 18^, resulting in enhanced metabolic activity and augmented tumourigenicity. Single-cell RNA expression profiling has identified a mitochondrial subtype in GB tumours that exhibits a mitochondrial signature and is dependent on mitochondrial oxidative phosphorylation (OXPHOS)^19^. Initially, it has been suggested that BTSCs rely heavily on OXPHOS to address their energy demands^20–22^. Later studies have reinforced this paradigm and suggested that at least subtypes of BTSCs including the slow-cycling rely on mitochondrial respiration^14^. This highlights that molecules that are able to affect mitochondrial respiration might be an interesting effective therapeutic strategy to eradicate BTSCs and suppress this devastating tumour.

Due to the dependency of multiple tumours on mitochondrial respiration^23–26^, targeting this metabolic pathway has emerged as an attractive anticancer strategy^27–29^. However, the translation of the currently available OXPHOS inhibitors into clinical practice has been hindered by poor potency, e.g., metformin and other biguanides^28, 30, 31^, or high toxicity, e.g., IACS-010759, oligomycin, rotenone, BAY 87-2243 and ASP4132^26, 29, 32, 33^. Here, we investigated the impact of mubritinib on BTSC fate and tumourigenesis. Mubritinib was initially described as an ERBB2 inhibitor, but recently found to inhibit OXPHOS in acute myloid leukemia model (AML)^34^. We report, for the first time, that mubritinib potently impairs stemness and growth of patient-derived BTSCs harboring different oncogenic mutations through targeting mitochondrial respiration. Importantly, we show that mubritinib is a brain penetrant drug that significantly suppresses GB tumour progression and sensitized their response to IR therapy. By performing profound toxicological and behavioral studies in preclinical animal models, we show that mubritinib has a well-tolerated and safe profile and holds promise for future clinical trials.

## Results

### Mubritinib is a potent inhibitor of OXPHOS in BTSCs

To determine the impact of mubritinib on the overall mitochondrial respiration in BTSCs, we examined oxygen consumption rate (OCR) as a measure of electron transport chain (ETC) activity. Multiple patient-derived BTSCs were treated with different concentrations of mubritinib and subjected to the ultra-sensitive real-time Resipher system (Lucid Scientific) or to the high-resolution respirometer, Oroboros, to measure the OCR (**Fig 1a-f**). These analyses revealed a significant and a dose-dependent reduction in basal mitochondrial respiration in mubritinib-treated BTSCs compared to vehicle control starting at very low concentrations in the range of 20 nM (**Fig. 1a-f**). Next, we evaluated the impact of mubritinib on the bioenergetics fitness of BTSCs by assessing the maximal mitochondrial and spare respiratory capacity and found a significant decrease in these respiration parameters upon mubritinib treatment, suggesting that mubritinib alters the mitochondrial reserve in BTSCs (**Fig. 1d-f**). Then, we asked if mubritinib inhibition of mitochondrial respiration in BTSCs is through impairment of complex I of the ETC as previously reported mechanism in AML^34^. To address this question, we performed rescue experiments using the Saccharomyces cerevisiae rotenone-insensitive NDI1 gene (**Fig. 1g**), and found that the ectopic expression of NDI1 gene in multiple BTSCs rescues the mubritinib-induced reduction in basal (**Fig. 1h-i**), maximal and spare respiratory capacity (**Fig. 1j-k**), suggesting that mubritinib impairs complex I activity to inhibit mitochondrial respiration in BTSCs.

**Figure 1.**
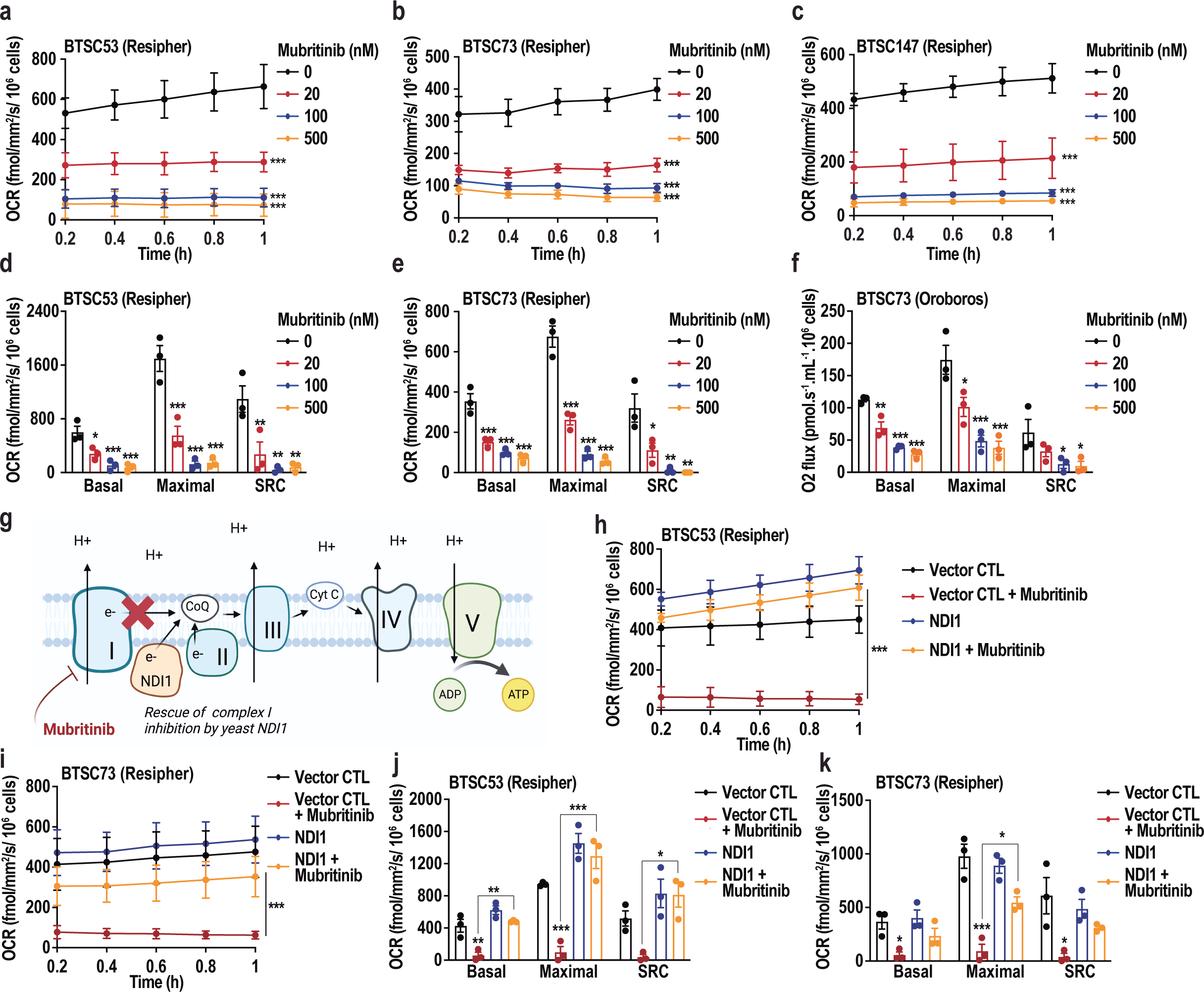
Mubritinib inhibits mitochondrial respiration through complex I. (**a-c**) Patient- derived BTSCs, BTSC53 (a), BTSC73 (b) and BTSC147 (c) were subjected to Resipher analysis to measure the basal oxygen consumption rate (OCR) following mubritinib treatment. (**d-e**) Maximal mitochondrial respiration and spare respiratory capacity (SRC) were measured in BTSC53 (d) and BTSC73 following mubritinib treatment using Resipher system (e). (**f**) Oxygene (O2) flux was measured using an Oroboros system. (**g**) Schematic diagram of the mitochondrial electron transport chain with ectopic NDI1 expression is presented. (**h-i**) BTSC53 (h) and BTSC73 (i) expressing vector control (CTL) or NDI1 were treated with vehicle control or 500 nM of mubritinib and subjected to Resipher analysis to measure the basal OCR. (**j-k**) Maximal respiration and SRC were measured using Resipher system in BTSC53- (j) and BTSC73- (k) expressing vector control (CTL) or NDI1 treated with either vehicle control or 500 nM mubritinib. Data are presented as the means ± SEM, n = 3. One-way ANOVA followed by Dunnett’s test vs vehicle CTL (a-f), one-way ANOVA followed by Tukey’s test (h-k). *p < 0.05, **p < 0.01, ***p < 0.001.

### Mubritinib impairs BTSC transcriptional state signatures and arrests their growth

Having established that mubritinib drastically impairs OXPHOS in BTSCs, and given the possible reliance of subsets of BTSC on OXPHOS, we thus asked if inhibition of OXPHOS by mubritinib impairs BTSC growth. We employed increasing concentrations of mubritinib (0.020 to 0.5 µM) on multiple patient-derived BTSC lines that harbour different genetic mutations and a murine BTSC line (mGB2) (**Fig. 2a, Supplementary** Fig. 1a and **Supplementary Table 1**). We observed a significant inhibition of BTSC growth in all the BTSCs tested at very low concentrations of mubritinib in nM range, following 4 days of treatment with mubritinib (**Fig. 2a**). Cell viability analysis revealed that BTSCs have different sensitivities to mubritinib treatment, we, therefore, asked whether it is correlated with specific genetic mutations in the patient-derived BTSCs (**Supplementary Table 1**). We observed that BTSCs harboring EGFR mutations (EGFRvIII or activating G598V mutation) are more sensitive to mubritinib, with a lower area under curve (AUC) compared to BTSCs lacking the EGFR mutations (**Fig. 2b**). However, TP53 mutation and methylguanine methyltransferase (MGMT) status did not impact mubritinib sensitivity (**Supplementary** Fig. 1b-c). Next, we investigated whether mubritinib sensitivity could be correlated with the basal OCR of BTSCs. We observed a positive correlation between the basal OCR and sensitivity to mubritinib (**Fig. 2c**), suggesting that the inhibitory effect of mubritinib on BTSC growth is due to OXPHOS impairment. To validate this mechanism, we assessed the impact of mubritinib on BTSCs growth following ectopic expression of the NDI1 gene. We found that NDI1 rescues the mubritinib-induced reduction in BTSC growth (**Fig. 2d-e**).

**Figure 2.**
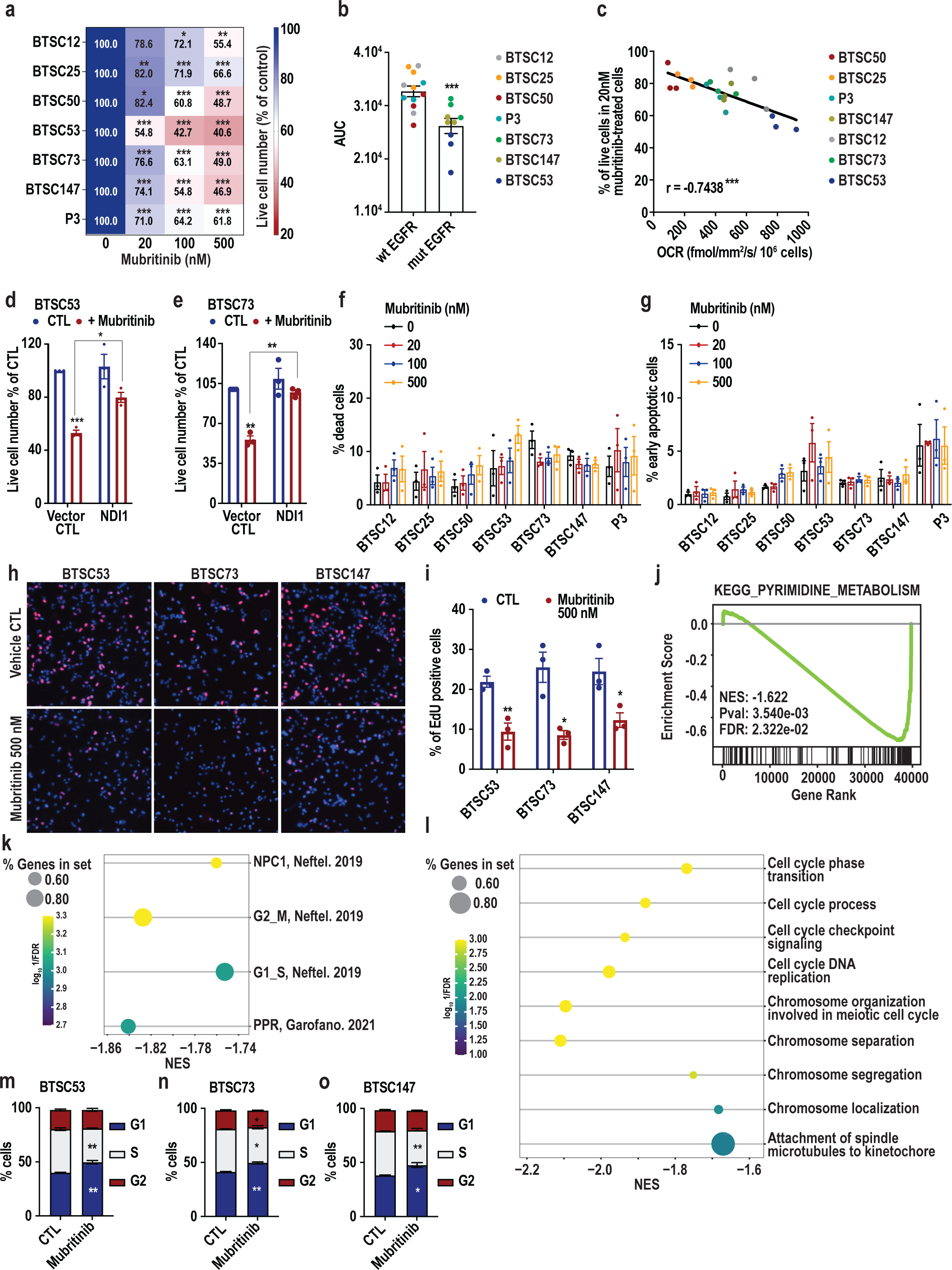
Mubritinib decreases proliferation of BTSCs without induction of cell death. (**a**) Multiple patient-derived BTSCs, BTSC12, BTSC25, BTSC50, BTSC53, BTSC73, BTSC147 and P3, were exposed to increasing concentrations of mubritinib (0 to 500 nM) for 4 days, following by live cell counting. (**b**) The area under the mubritinib dose-response curves (AUC) of wild-type EGFR (wt) BTSCs (#12, #25, #50, P3) and mutated (mut) EGFR BTSCs (#53, #73 and #147) were plotted. (**c**) Pearson correlation analysis of basal oxygen consumption rate (OCR) and the percentage of live cells in BTSC following 4 days treatment with 20 nM of mubritinib was performed. (**d-e**) The number of live cells in BTSC53- (d) and BTSC73 (e) - expressing vector control or NDI1 following 4 days of 500 nM mubritinib treatment was measured. (**f-g**) The percentages of dead cells (PI positive) and early apoptotic cells (Annexin V positive and PI negative) were measured by Annexin V/PI double staining followed by flow cytometry in BTSCs treated with increasing concentrations of mubritinib for 4 days. (**h-i**) EdU incorporation was analyzed by immunofluorescence imaging in BTSCs (#53, #73 and #147) after 4 days of 500 nM mubritinib treatment. Representative images of EdU (red) staining are shown (h). Nuclei were stained with DAPI (blue). The number of EdU positive cells was quantified with Fiji software (i). (**j-l**) Gene set enrichment analysis of deregulated genes in BTSC147 treated with 500 nM of mubritinib for 24 h demonstrates enrichment for gene sets corresponding to pyrimidine metabolism (**j**), NPC1, G2M and G1S states defined by Neftel et al.; the proliferative (PPR) signature defined by Garofano et al. (k) and cell-cycle related pathways (l). (**m-o**) Cell cycle distribution was assessed by flow cytometry after PI staining in BTSC53 (m), BTSC73 (n) and BTSC147 (o) following 24 h of treatment with 500 nM of mubritinib. Data are presented as the means ± SEM, n = 3. One-way ANOVA followed by Dunnett’s test vs vehicle CTL (a), unpaired two-tailed t test (b, i, m-o), one-way ANOVA followed by Tukey’s test (d, e). *p < 0.05, **p < 0.01, ***p < 0.001.

We next asked if the decrease in cell number in response to mubritinib is due to inhibition of proliferation or induction of cell death. Annexin V/propidium iodide (PI) flow cytometry analysis revealed that there is no significant induction of cell death or early apoptosis in any of the BTSCs tested even at concentrations up to 10 µM of mubritinib (**Fig. 2f-g and Supplementary** Fig. 1d). On the other hand, by performing 5-ethynyl-2′-deoxyuridine (EdU) assay, we observed a drastic reduction in the number of EdU-incorporating BTSCs in mubritinib-treated conditions (#53, #73 and #147) (**Fig 2h-i**). These results confirm that the decrease in cell number following mubritinib treatment is not due to cell death but it is likely attributed to inhibition of proliferation. To gain further mechanistic insights on changes induced by mubritinib in BTSCs, we performed RNA-seq analysis 24 h following mubritinib treatment (**Supplementary** Fig. 2a-b). Interestingly, mubritinib resulted in downregulation of pyrimidine metabolism pathway, and alanine, aspartate and glutamate metabolism-related pathways (**Fig. 2j and Supplementary** Fig. 2c). These metabolic pathways are tightly associated with ETC activity^35, 36^. In support to the proliferation inhibition observed by mubritinib, RNAseq analysis revealed a significant downregulation of the proliferative (PPR) signature described by Garofano et al., ^19^ and is enriched in pathways associated with cell cycle progression, DNA replication and mitosis (**Fig. 2k**). Furthermore, analysis of Neftel et al.,^37^ cell state signatures of GB, revealed that mubritinib significantly downregulates the neural-progenitor-like cell (NPC1) state and the proliferative G1/S and G2/M states (**Fig. 2k**). GOBP enrichment analysis showed downregulation of cell-cycle related pathways such as cell cycle DNA replication and cell cycle checkpoint signaling (**Fig. 2l**). To validate the role of mubritinib on cell cycle deregulation, we performed functional cell cycle profiling using flow cytometric quantitation of DNA content by PI. A significant accumulation of cells in the G1 phase and a decrease in the percentage of cells in S phase were observed in mubritinib treated-BTSCs (**Fig. 2m-o**). Altogether, these data support that mubritinib alters mitochondrial activity to impair cycling of BTSCs.

### Mubritinib impairs self-renewal and results in downregulation of stemness-related pathways

It has been suggested that OXPHOS activity maintains GSCs stemness^21, 38, 39^. By analysis of RNAseq data, we observed that mubritinib downregulates the NPC state and the PPR signature that are enriched in markers of neural stem/progenitor cells. We thus asked if the impairment of mitochondrial respiration by mubritinib impacts BTSC self-renewal. To address this question, we subjected three patient-derived BTSC lines (#53, #73, #147) and the murine BTSC line (mGB2) to extreme limiting dilution assays (ELDA) in the absence and presence of mubritinib. ELDA analysis showed that mubritinib treatment robustly attenuated self-renewal in all the BTSC lines tested (**Fig. 3a-c and Supplementary** Fig. 3a). To further assess the impact of mubritinib on BTSC stemness, we assessed the levels of the different stemness markers in multiple BTSCs by immunoblotting. Our analysis revealed a strong reduction in the protein levels of the tested stemness markers i.e. SOX2, Olig2, Nestin and active (cleaved) Notch1 in BTSCs treated with mubritinib compared to control (**Fig. 3d-f and Supplementary** Fig. 3b). Strikingly, immunoblotting analysis revealed that mubritinib treatment has no significant impact on the levels of stemness markers in non-oncogenic NSCs (**Supplementary** Fig. 3c). Altogether, our data suggest that mubritinib impairs self-renewal and stemness-related pathways specifically in BTSCs.

**Figure 3.**
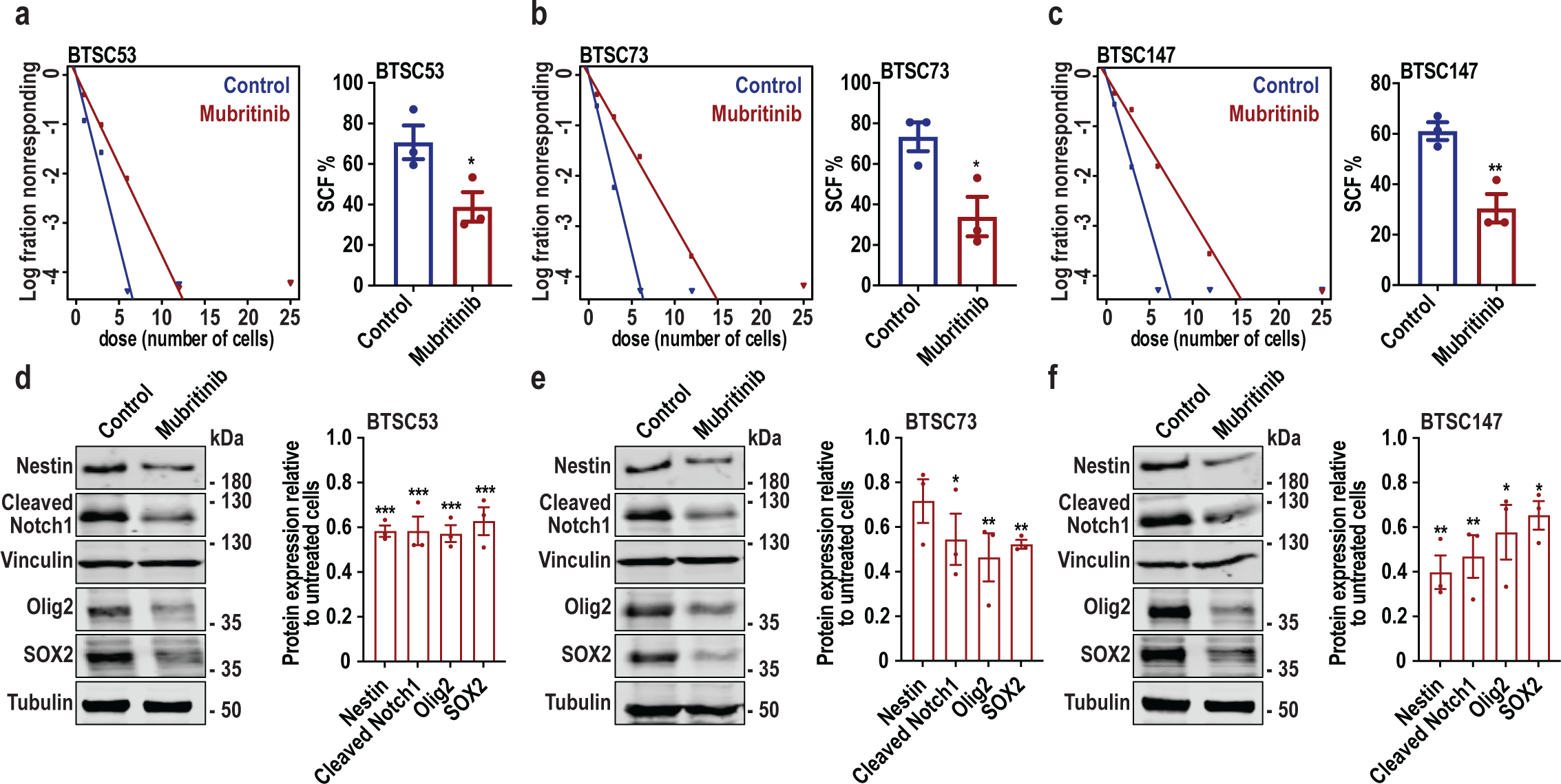
Mubritinib impairs BTSC stemness properties. (a-c) BTSC53 (a), BTSC73 (b) and BTSC147 (c) were subjected to extreme limiting dilution analysis to estimate the stem cell frequencies (SFC) following treatment with 500 nM mubritinib or vehicle control. (**d-f**) BTSC53 (d), BTSC73 (e) and BTSC147 (f) were treated for 4 days with 500 nM of mubritinib or vehicle control and subjected to immunoblotting using the antibodies indicated on the blots. Vinculin and Tubulin were used as loading control. Densitometric quantification of Nestin, cleaved Notch1, Olig2 and SOX2 protein level normalized to their corresponding loading control are presented. Data are presented as the mean ± SEM, n = 3. Unpaired two-tailed t test (a-c). One-way ANOVA followed by Dunnett’s vs vehicle CTL, *p < 0.05, **p < 0.01, ***p < 0.001.

### Mubritinib is a brain penetrant drug that impairs tumourigenesis in patient-derived BTSC xenografts and syngeneic models

We next set out to investigate the functional relevance of these findings to GB *in vivo*. Since insufficient drug exposure in the brain is one of the major hurdles in treatment of brain tumours, we thus asked whether mubritinib could cross the blood-brain barrier (BBB). To address this question, we designed pharmacokinetic experiments to assess the bioavailability of mubritinib in the blood and the brain. After single intraperitoneal injection with mubritinib, animals were sacrificed at different time points, plasma and brain samples were collected and subjected to mubritinib quantification by liquid chromatography / tandem mass spectrometry analysis (LC/MS/MS) (**Fig. 4a**). Striking, we found that mubritinib accumulates in the brain, at a brain/blood ratio of 2 to 3.4-fold after 12 h to 36 h, respectively (**Fig. 4b**). It reaches a Cmax of 2252 ng/mL after 4 h in the plasma, and 4325 ng/mL after 24 h in the brain (**Fig. 4b**). Importantly, after 36 h mubritinib was still detectable in both plasma and brain (**Fig. 4b**). These data highlight that mubritinib bypasses the BBB, accumulates in the brain and cleared slowly from the brain. Having established that mubritinib penetrates through the BBB, we therefore, investigated whether mubritinib could suppress brain tumours. To begin with, we intracranially implanted the murine-derived BTSCs (mGB2) into the brains of immunocompetent C57BL/6N mice (**Fig. 4c**). The animals were randomized to receive either mubritinib (6 mg/kg) or vehicle control intraperitoneally (**Fig. 4c**). Tumour formation was monitored by bioluminescent imaging of luciferase signal using an *in vivo* real-time optical imaging system (Biospace) (**Fig. 4d-e**). Luciferase imaging revealed a marked decrease in brain tumour growth in mubritinib-treated mice compared to vehicle treated-mice (**Fig 4d-e**). At ∼40 days following surgery, mice receiving vehicle control formed malignant brain tumours and were at endpoints as assessed by major weight loss and neurological signs. Strikingly, mubritinib delayed mGB2 xenograft tumourigenesis and significantly extended the lifespan of animals (**Fig. 4f**). To confirm the effect of mubritinib treatment on GB tumour growth and animal survival, we performed parallel experiments with a patient-derived line (BTSC147), as described for mGB2. 100 days following intracranial implantation of BTSC147 in RAGγ2C^−/−^ mice, the group receiving vehicle-control formed malignant brain tumours while mice treated with mubritinib exhibited a significant delay in tumour formation with a significant ∼3-fold decrease of luciferase activity (**Fig. 4g-h**). Importantly, mubritinib significantly extended the lifespan of animals bearing BTSC147- tumours (**Fig. 4i**). Using another patient-derived BTSC model (BTSC73), we validated the effect of mubritinib in expanding animal lifespan (**Fig. 4j**). Having established that mubritinib impairs stemness and self-renewal in BTSCs *in vitro,* we next asked whether it alters stemness in brain tumours *in vivo*. To address this question, we orthotopically inoculated BTSC73 into 8-week RAGγ2C^−/−^ mice. After 7 days, mice were randomized into two treatment arms: vehicle control or mubritinib (6 mg/kg). After 30 days of surgeries, all mice were sacrificed, brains were sectioned and stained with Olig2 and SOX2. Strikingly, we observed a strong reduction in the number of Olig2 and SOX2 positive cells in mubritinib treated-mice (**Fig. 4k-l**). Altogether, these data demonstrate that mubritinib crosses the BBB, effectively delays GB growth, decreased GB stemness and conferred a survival advantage.

**Figure 4.**
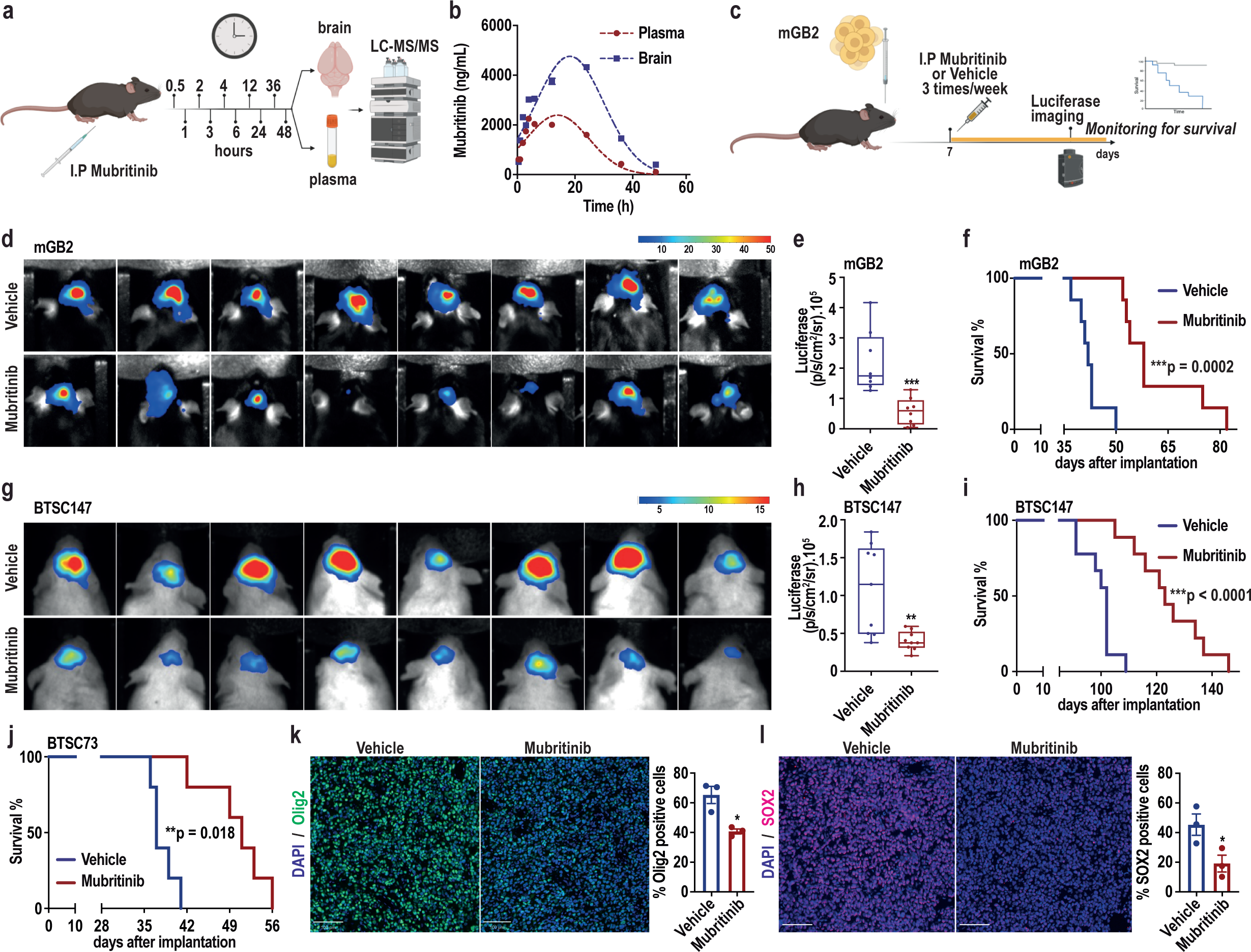
Mubritinib, a brain penetrant drug, delays BTSCs tumourigenesis. (a-b) Schematic diagram of the pharmacokinetic analysis is shown (a). C57BL/B6N mice were treated by I.P with a single dose of mubritinib (6 mg/kg). Brain and plasma were collected at different time point (0.5 h to 48 h) after injection. Mubritinib was quantified by LC-MS/MS (b). (**c**) Schematic diagram of the experimental procedure in which luciferase-expressing mGB2 cells (500K) were intracranially injected into C57BL/B6N mice. One week after implantation mice were randomized in 2 groups: vehicle control or mubritinib (6 mg/kg). Mice were treated 3 times per week (Monday, Wednesday and Friday). Luciferase imaging was used to follow tumour progression. (**d-e**) Bioluminescence images (d) and quantification of luciferase activity (e) 6 weeks after implantation are presented. n = 8 mice. (**f**) Kaplan–Meier (KM) survival plot was graphed to evaluate mice lifespan in each group, mice were collected at end stage (n = 7 mice). (**g-i**) Luciferase-expressing BTSC147 were intracranially injected into RAGγ2C^−/−^ mice. Bioluminescence images (g) and quantification of luciferase activity (h) 9 weeks after implantation are presented (n = 9 mice). KM survival plot was graphed to assess animal lifespan (n = 9 mice) (i). (**j**) BTSC73 were intracranially injected into RAGγ2C^−/−^ mice and KM survival plot was graphed to assess animal lifespan between vehicle and mubritinib treated mice (n = 5 mice). (**k-l**) Representative immunofluorescence images and the quantification of Olig2 (green) (k) and SOX2 (red) (l) positive cells in BTSC73 intracranial xenografts in mice treated with mubritinib or vehicle are shown. Nuclei were stained with DAPI (blue). The number of Olig2 and SOX2 positive cells was quantified with Fiji software (n = 3 mice). Scale bar = 100 µm. Data are presented as the mean ± SEM. Unpaired two-tailed t test (e, h, k and l), log-rank test (f, i and j). **p < 0.01, ***p < 0.001.

### Targeting OXPHOS by mubritinib sensitizes GB tumours to IR

Recent proteomic analysis of GB tumours revealed an enrichment of OXPHOS-associated proteins in recurrent compared to primary GB tumours, suggesting that OXPHOS could be involved in GB treatment resistance^40^. Furthermore, our RNAseq analysis showed that mubritinib downregulates homologous recombination related-pathway, a major pathway for DNA repair that is involved in BTSC radioresistance (**Fig. 5a**)^41, 42^. We thus asked whether BTSCs upregulate the mitochondrial respiration in response to IR-induced insults and DNA damage, and if inhibition of OXPHOS by mubritinib could improve GB response to IR. To address these questions, we first exposed multiple patient-derived BTSCs to either 2 Gy or 4 Gy of IR followed by OCR measurement via Resipher. We observed an increase in OCR in irradiated BTSCs compared to control cells (**Fig. 5b-c**). Next, we transplanted luciferase expressing-BTSC73 into the brain of immunodeficient mice. Mice were randomized to receive one of the four treatment arms: vehicle control, mubritinib (6 mg/kg), irradiation (2 cycles of 2 Gy) or a combination of mubritinib and IR (**Fig. 5d**). Luciferase imaging, at 21 days following surgeries, showed a delay in tumour formation in the group receiving either mubritinib or IR monotherapy, with a ∼5-fold decrease in luciferase activity compared to control group. Interestingly, a much higher reduction (∼14-fold) in luciferase activity was observed in mice receiving a combinational therapy of mubritinib and IR (**Fig. 5e-f**). The tumour volume was estimated by H&E staining 30 days after surgery. As observed by luciferase imaging, while IR alone and mubritinib alone decreased tumour volume compared to control, the combination of IR and mubritinib resulted in smaller tumours (**Fig. 5g-h**). Importantly, KM survival plots revealed that animals receiving the combination of IR and mubritinib had survival advantage over the animals receiving either IR or mubritinib monotherapy (**Fig. 5i-j**). Indeed, a ∼ 50% extension of the survival rate was observed in mice receiving IR and mubritinib compared to the control group (**Fig. 5i-j**). To further validate these results, we implanted a second patient-derived BTSCs (#53) into the brains of RAGγ2C^−/−^ mice. The medians of survival of mice were 57, 70 and 67 days in vehicle control, mubritinib and irradiated mice respectively. Interestingly, the median survival was extended to 84 days in mice receiving the combination of IR and mubritinib (**Fig. 5k-l**). Overall, these results suggest that combinational therapy of mubritinib and IR represents a promising strategy to treat GB tumours.

**Figure 5.**
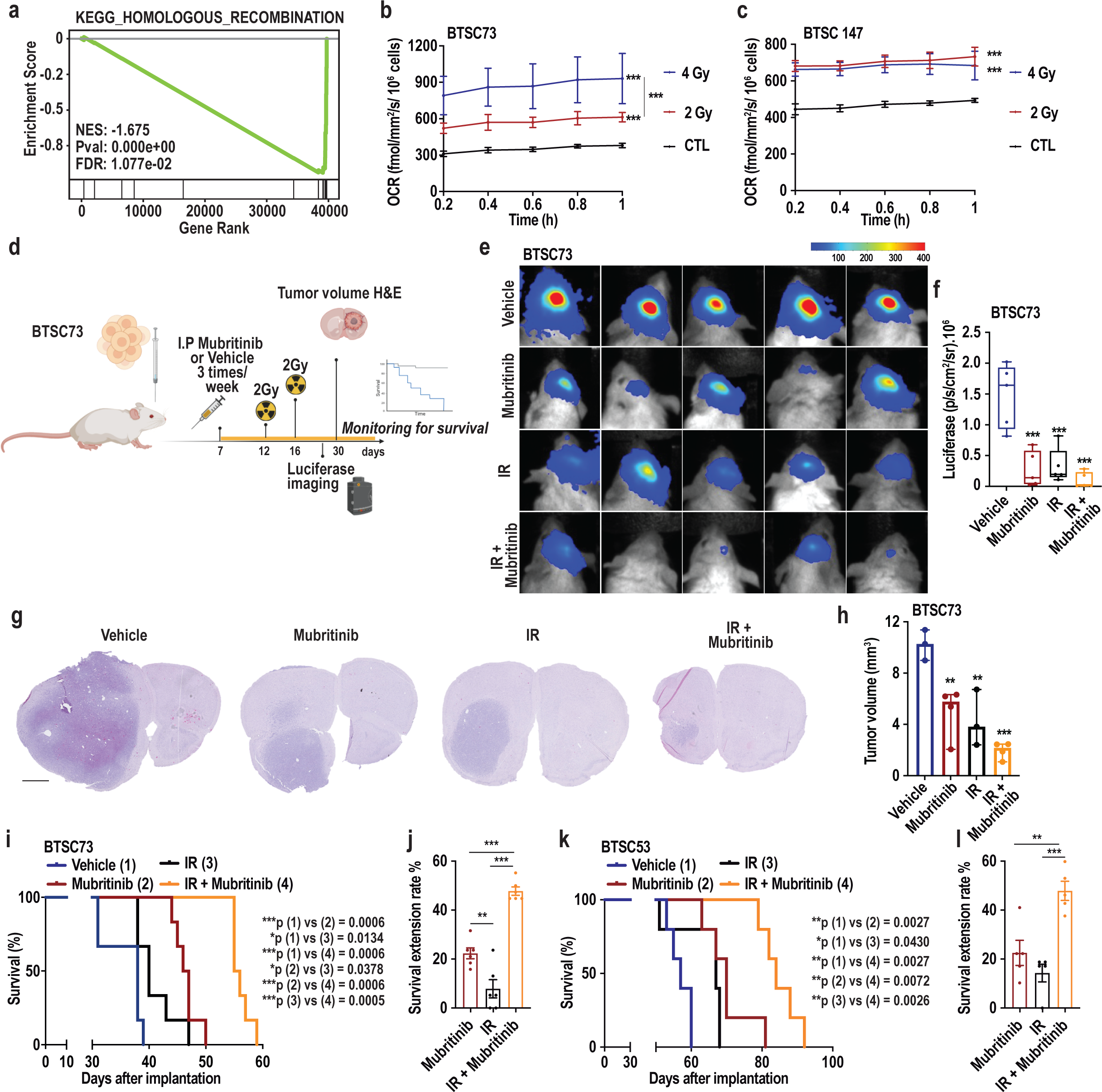
Mubritinib sensitizes GB tumour to ionizing radiation. (**a**) Gene set enrichment analysis of deregulated genes in BTSC147 treated with 500 nM of mubritinib for 24 h demonstrates enrichment for gene sets corresponding to homologous recombination. (**b-c**) 2 Gy or 4 Gy irradiated BTSC73 (b) and BTSC147 (c) were subjected to Resipher analysis to measure the basal oxygen consumption rate (OCR) 16 h following irradiation. (**d**) Schematic diagram of the experimental procedure in which luciferase-expressing BTSC73 cells (300K) were intracranially injected into RAGγ2C^−/−^ mice. One week after implantation mice were randomized in 2 groups: vehicle control or mubritinib (6 mg/kg). Mice were treated 3 times per week (Monday, Wednesday and Friday). At day 12, mice from the 2 groups were randomized to receive or not IR (2 Gy). (**e-f**) Bioluminescence images (e) and quantification of luciferase activity (f) 3 weeks after implantation are presented. n = 5 mice. (**g-h**) H&E stainings were performed at day 30 (g). Scale bar, 1mm. Tumour volume was estimated based on H&E stainings (h). (**i**) Kaplan–Meier (KM) survival plot was graphed to evaluate mice lifespan in each group, mice were collected at end stage (n = 6 mice). (**j**) Survival extensions of mice bearing BTSC73-derived tumours treated with mubritinib, IR, or mubritinib + IR relative to those treated with the vehicle control were calculated. (**k-l**) BTSC53 were intracranially injected into RAGγ2C^−/−^ mice. KM was plotted to evaluate mice lifespan in each group (k), and survival extensions were calculated (l) (n = 5 mice). Data are presented as the mean ± SEM. One-way ANOVA followed by Tukey’s test (b, c, f, h, j and l) and log-rank test (i and k). **p < 0.01, ***p < 0.001.

### Mubritinib doesn’t induce damage to healthy cells and has a well-tolerated profile

Having established that mubritinib impairs BTSC growth and tumourigenesis *in vitro* and *in vivo*, raised the questions of the impact of mubritinib on non-oncogenic normal cells. To address this question, we employed wild type human normal neural progenitor cells (NPCs), human hepatocytes (HepaSH) and murine post-mitotic primary mouse neurons (**Fig. 6a**). First, we assessed cell proliferation and viability in hNPCs, *in vitro*, following mubritinib treatment by using Calcein-AM/DAPI double staining to stain live and dead cells respectively (**Fig. 6b**). Interestingly, at concentrations where mubritinib causes inhibition of proliferation in BTSCs, no difference in number of live hNPCs was observed (**Fig. 6b-c**). Furthermore, by assessing the number of dead cells (DAPI-positive), we observed that mubritinib did not induce cell death in hNPCs (**Fig. 6b and d**). Next, we assessed the toxicity of mubritinib on hepatocytes and neurons. Phase-contrast examinations showed no morphological alterations in mubritinib-treated human hepatocytes compared to the hepatotoxic drug Bosentan (**Fig. 6e**). In addition, the MTT cell viability assays, revealed no significant decline in the viability of HepaSH (**Fig. 6f**) or mouse neurons (**Fig. 6g**) following mubritinib treatment.

**Figure 6.**
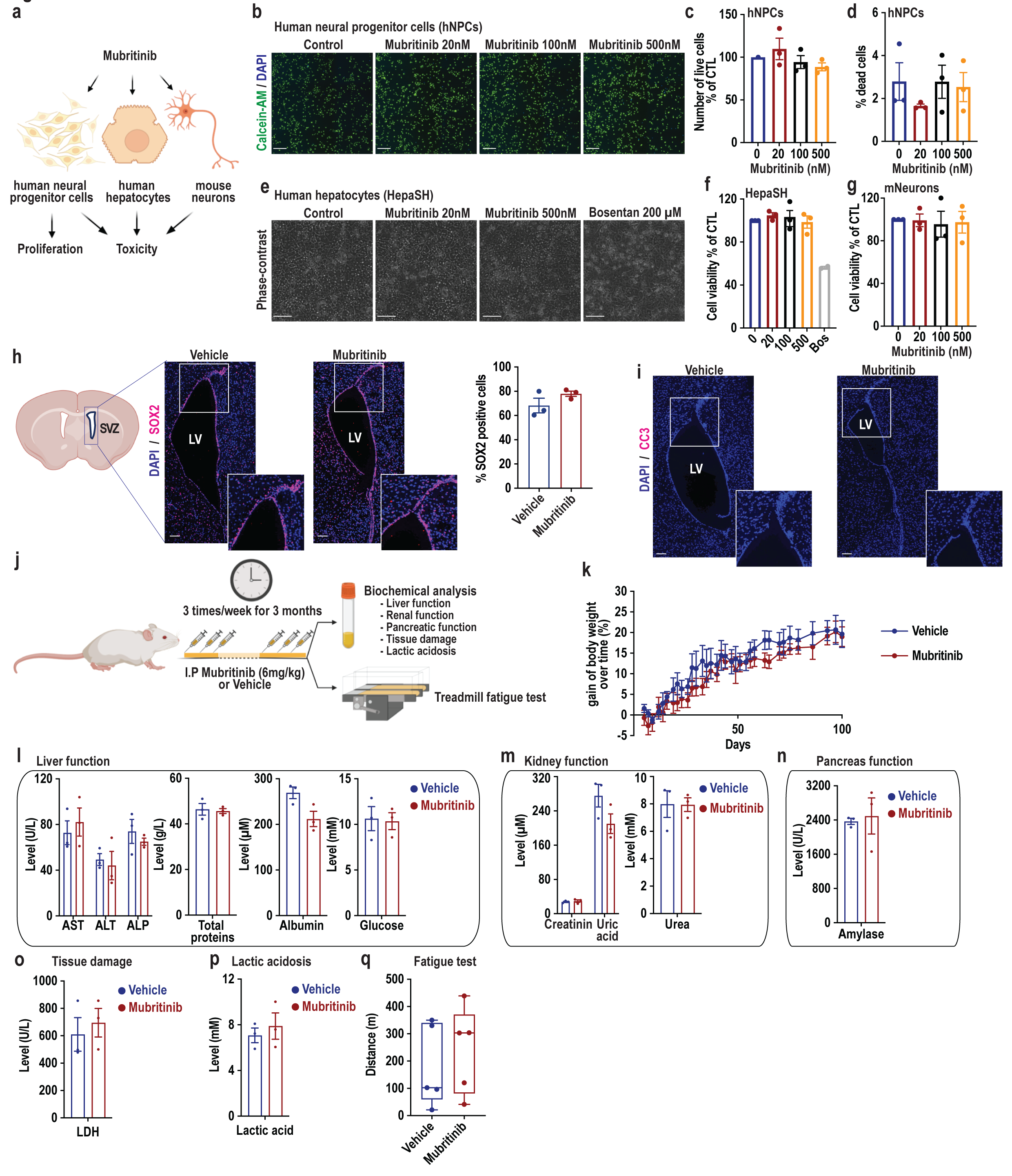
Mubritinib does not impact normal non-oncogenic cells and has a safe profile *in vivo*. (**a**) Schematic diagram of the *in vitro* analysis of mubritinib on normal cells. (**b-d**) Human neural progenitor cell line (hNPC) was treated with increasing concentration of mubritinib (0 to 500 nM) for 4 days and subjected to Calcein-AM/DAPI double staining. Representative images are shown (b). Scale bars = 100 µm. The number of live cells (calcein-am positive) (c) and dead cells (DAPI positive) (d) were quantified by Fiji software. (**e-f**) The human hepatocytes, HepaSH, were treated for 4 days with mubritinib or the hepatotoxic drug, Bosentan. Phase-contrast images were captured after 4 days (e). Scale bars = 100 µm. Cytotoxicity was measured by MTT after 4 days (f). (**g**) Cell viability of mouse cortical neurons (mNeurons) was measured by MTT after 4 days of mubritinib treatment. (**h-i**) Representative immunofluorescence images of SOX2 (h) and cleaved caspase 3 (CC3) (i) in the SVZ of mouse brains with indicated treatments are shown (n = 3 mice). Scale bars = 100 µm. The number of SOX2 positive cells was quantified with Fiji software (n = 3 mice). (**j**) Schematic diagram of the *in vivo* toxicological studies is presented. Biochemical analysis and treadmill fatigue test were performed on RAGγ2C^−/−^ mice treated for 3 months with vehicle control or mubritinib (6 mg/kg), 3 times per week (Monday, Wednesday and Friday). (**k**) Body weight of mice is presented (n = 5 mice). (**l-p**) Biochemical analyses to evaluate liver (l), kidney (m), pancreatic function (n), tissue damage (o) and lactate acidosis (p) were performed (n = 3 mice). (**o**) Treadmill fatigue test was performed and the distances ran by mubritinib-treated group and vehicle control were measured (n = 5 mice). Data are presented as the mean ± SEM.

Induction of toxicity in normal brain cells have been reported with many chemotherapeutic agents. For instance, the anti-GB drug carmustine, has been shown to impact normal brain cells, and increase the cell death in the lateral subventricular zone (SVZ) where the NPCs reside^43^. We first assessed whether mubritinib treatment alters the healthy NPCs *in vivo*. Strikingly, we found that mubritinib had no apparent impact on the percentage of SOX2 positive cells NPCs (**Fig. 6h**). Furthermore, we performed immunostaining of cleaved caspase-3 (CC3) on brains from mubritinib-treated mice, and found no induction in apoptosis compared to vehicle-treated mice (**Fig. 6i)**. Altogether, the *in vitro* and *in vivo* data show that mubritinib impairs BTSCs without causing significant damages to healthy cells.

Next, we assessed the tolerability of mubritinib *in vivo*. We evaluated the systemic toxicity of mubritinib after 3 months (3 times/week) of repeated mubritinib treatment (6 mg/kg), by performing blood biochemical analysis and behavioral studies on mubritinib-treated mice (**Fig. 6j**). Assessment of body weight showed that mubritinib was well-tolerated at the tested dose without causing any loss of mice body weight (**Fig. 6k**). Through the blood biochemical analysis, we looked at liver function by measuring alanine amino transferase (ALT), aspartate amino transferase (AST), alkaline phosphatase (ALP), albumin, total proteins and glucose levels (**Fig. 6l**); kidneys function by measuring creatinin, urea and uric acid (**Fig. 6m**); and pancreas function by measuring amylase (**Fig. 6n**). Strikingly, no significant changes in the levels of these markers have been observed in mubritinib-treated mice compared to vehicle control. In addition, no difference in LDH level has been observed, indicating no induction of any tissue damage following mubritinib treatment (**Fig. 6o**). Lactic acidosis is a frequently reported side effect and a major concern that leads to the termination of many clinical trials of OXPHOS inhibitors^29^, we thus measured the level of lactic acid. Strikingly, mubritinib did not induce increase in the levels of lactic acid (**Fig. 6p**). Next, we performed a non-voluntary preclinical assay of fatigue-like behavior, the treadmill fatigue test, on mice following 3 months of repeated mubritinib treatment (**Fig. 6i**). After 2 sessions of training, the fatigue-test was performed. Interestingly, we did not detect any fatigue-like behavior in mubritinib-treated mice compared to control. In fact, the distances ran by the 2 groups of mice were the same (**Fig. 6q**). Taken together, these data indicated that at anti-GB effective doses, mubritinib has a well-tolerated and safe profile. It could represent a promising safe therapeutic agent for the treatment of GB tumours.

## Discussion

GB is the most aggressive and deadly brain tumour with dismal prognosis despite multimodal therapies^44^. GB tumours exhibit remarkable cellular heterogeneity, with a population of BTSCs at the apex of the differentiation hierarchy^45^. BTSCs display potent tumour-initiating capacity, and promote malignant behaviors associated with disease progression and relapse^45–47^. Therefore, targeting BTSCs is crucial for improving GB treatment and overcoming therapeutic resistance. In the present work, we established, for the first time, that mubritinib is a promising drug to target cancer stem cells in GB. We employed murine-derived and multiple patient-derived BTSCs with different genetic background, and found that mubritinib suppresses BTSC stemness pathways, self-renewal and proliferation. Mechanistically, by employing mitochondria respiration assays and rescue experiments using the ectopic expression of the Saccharomyces cerevisiae NDI1 gene, we provide compelling evidence that mubritinib acts on complex I of the ETC to impair mitochondrial respiration and BTSC growth. By performing pharmacokinetics, we established that mubritinib is a brain-penetrant drug. Using patient-derived BTSC orthotopic xenografts and syngeneic murine GB preclinical models, we have established that mubritinib delays GB tumourigenesis, sensitizes GB response to IR and expands lifespan of GB tumour-bearing animals. Importantly, biochemical, toxicological and behavioral studies showed that mubritinib exhibits a safe profile. Altogether, these data unravel that mubritinib is a potent drug to impair BTSCs and suppress GB tumourigenesis as a stand- alone therapy or in combination with IR.

Mubritinib, also known as TAK165, is a small molecule that was developed in the early 2000s and has been initially reported as a specific ERBB2 inhibitor^48^. However, later studies demonstrated that gastric cancer cells that exhibit dramatic reduction in expression of various tyrosine kinase receptors (TKR), including HER2/ERBB2, are resistant to TKR inhibitors but sensitive to mubritinib. This suggests that mubritinib acts via alternative mechanisms that do not involve inhibition of HER2/ERBB2^49^. In a more recent work, Baccelli and colleagues have demonstrated that mubritinib selectively inhibits growth of a subset of AML cells that rely on OXPHOS by inhibiting complex I of the ETC^34^.

Prior to our study, the impact of mubritinib on BTSCs and GB tumourigenesis was not investigated. Here, we provide solid evidence that mubritinib alters BTSC growth, stemness and tumourigenesis through impairment of complex I and subsequently inhibition of OXPHOS. Interestingly, we tested the effect of mubritinib on a panel of patient-derived BTSCs and we found a higher sensitivity of EGFR-mutated BTSCs. EGFR alterations have been observed in nearly 41-57% of GB^50–54^ and have been reported to induce an aggressive feature of GB and worse prognosis in certain studies^53^. Though EGFR mutated BTSCs were more sensitive, the EGFR, wild type BTSCs responded to mubritinib treatment suggesting that mubritinib could be used on a wide range of GB patients. An important feature of BTSCs is their metabolic plasticity, which allows them to shift between different metabolic pathways to survive nutrient deprivation or metabolic stress^22, 55–58^. However, certain genetic mutations impair this metabolic fitness of BTSCs and renders them more sensitive to OXPHOS inhibitors. For instance, homozygous deletion of the key glycolytic enzyme enolase 1 that occurs in 3.3% of GB patients, showed reduced capacity for compensatory glycolysis and increased sensitivity to the complex I inhibitor IACS-010759^59^. The oncogenic gene fusions of FGFR3-TACC3 (F3-T3) was described in 3% of human GB cases^60^ and is shown to activate OXPHOS and mitochondrial biogenesis and confers sensitivity to inhibitors of oxidative metabolism^61^. This suggests that mubritinib could be more effective on GBs harboring these genetic alterations.

Radioresistance has emerged as one of the major obstacles in GB therapy. BTSCs are central in conferring resistance to therapy and have been shown to be more radio-resistant than their non-stem cell counterparts in GB^45, 46, 62^. We previously showed that the increase in cellular respiration and spare respiratory capacity in BTSCs confer resistance to IR therapy^39^. In line with these studies, we show that BTSCs upregulate OXPHOS in response to IR, suggesting that BTSCs employ mitochondrial respiration to resist the IR insult. In radiology, targeting mitochondrial respiration is an emerging strategy to overcome radioresistance in the tumour hypoxic regions. OXPHOS inhibition results in a reduction in oxygen consumption in the well-oxygenated tumour, and therefore, an increase oxygen available for diffusion into hypoxic regions to improve radiation response^63, 64^. Repurposing metformin as cancer treatment is already being tested in a range of clinical trials for a variety of cancers, and it’s combination with TMZ and IR is being investigated as a therapeutic avenue for GB patients^65^. However, it did not confer a clinical benefit in patients with recurrent or refractory GB^66^. Furthermore, combination of metformin with chemoradiation did not improve overall or progression-free survival of patients with Non–Small Cell Lung Cancer^67^. Given the low potency of metformin as an OXPHOS inhibitor, the outcome of these clinical trials is not surprising. Our findings support that the use of mubritinib in combination with IR may provide a promising new avenue for better treatment of GB patients.

Drugs targeting mitochondria are emerging as promising antitumour therapeutics in preclinical models. However, in the last decades clinical development of agents targeting metabolism has proven challenging. A few clinical trials on OXPHOS inhibitors such as BAY87-2243, ASP4132 and IACS-010759, has been terminated due to dose-limiting toxicities and narrow therapeutic window of these agents^26, 28, 29^. Other clinical trials have failed due to the lack of anti-tumour effects^68^. In a very recent clinical trial, IACS-010759, was tested in patients with advanced solid tumours and AML^29^. Patients who were administered the drug developed unacceptable toxicities, including elevated blood lactate and peripheral neuropathy resulting in termination of the clinical trial^29^. At lower doses, this drug was not effective. Importantly, Yap et al., showed that IACS-010759 induces behavioral and physiological changes indicative of peripheral neuropathy in mouse models^29^, suggesting that preclinical models could predict these drug-induced injuries. In light of the outcome of these clinical trials, efforts should be directed towards thorough early evaluation of the safety of anticancer agents in preclinical context. Strikingly, our data showed that chronic mubritinib administration does not induce lactic acidosis or any significant changes in the biochemical serum assays. Furthermore, chronic mubritinib treatment did not induce behavioral changes or fatigue in the animals. Interestingly, the important accumulation of mubritinib that we observed in the brain suggests its efficacy at low doses, and therefore minimized side effects. Therefore, our data with mubritinib purport the recent bleak view of targeting mitochondrial respiration as an anti-cancer strategy, and encourages to further thoroughly evaluate the pharmacokinetics, pharmacodynamics and safety profiles of this drug.

In summary, our study identifies mubritinib as brain-penetrant drug with a safe profile and strong anti-GB activity in preclinical models. Given that mubritinib is already completed a phase I clinical trial in the context of ERBB2+ solid tumours, our work encourages and warrants future investigations for clinical translation and repurposing mubritinib for better management of GB tumours.

## Materials and methods

### Animals

All animal experiments were conducted under the institutional guidelines and were approved by the local ethics committee (agreement number: C335222). Housing room temperature and relative humidity were adjusted to 22.0 ± 2.0 °C and 55.0 ± 10.0%, respectively. The light/dark cycle was adjusted to 12 h lights-on and 12 h lights-off. Autoclaved water and irradiated food pellets were given ad libitum.

### Patient-derived and murine BTSC cultures

The human BTSCs 12, 25, 50, 53, 73, and 147 were generated from patient GB tumours by Dr. Samuel Weiss team at the University of Calgary^69, 70^. The P3 line was derived from a GB tumour as previously described^71^. Cells were characterized for major mutations in GB including EGFRvIII, p53, PTEN, and IDH1 status (**Supplementary Table 1**). The murine GB cells, mGB2, were kindly provided by Dr. Peter Angel^72^. Prior to use, BTSCs were recovered from cryopreservation in 10% DMSO and cultured in low attachment flasks as spheres in DMEM/F12 media (Fisher Scientific, #11530566) supplemented with 2 mM GlutaMAX (ThermoFischer, #35050061), 1X B-27 supplement (ThermoFischer, #12587010), 2 µg/ml heparin (Sigma-Aldrich, #H3149), 20 ng/ml EGF (Peprotech, #AF-100-15), 20 ng/ml bFGF (Peprotech, #AF-100-18B), 100 U/ml penicillin-streptomycin (Fisher Scientific, #11548876). All cell lines were tested negative for mycoplasma. All cells were kept in a humidified 5% CO2 incubator at 37°C.

### Mouse NSC cultures

Primary adult mouse NSC were isolated from the subventricular zone (SVZ) of 6 week-old C57BL/6N mouse brain, expanded *in vitro* and grown as neurospheres in low attachment flasks in DMEM/Ham’s F-12 medium supplemented with 1X B-27 (ThermoFischer, #12587010), 20 ng/mL mEGF (Cell Signaling, #5331SC) and 20 ng/mL mFGF (Abbiotec, #600182)^73^.

### ReNcell CX human neural progenitor cell cultures

The human NPC, ReNcell CX line, derived from the cortical region of human fetal brain tissue (Millipore, #SCC007) were cultured as previously described^74^. The ReNcell CX cells were cultured on matrigel-coated plates or flasks and maintained in DMEM/F12 media (Fisher Scientific, #11530566) supplemented with 2 mM GlutaMAX (ThermoFischer, #35050061), 1X B-27 supplement (ThermoFischer, #12587010), 2 µg/ml heparin (Sigma-Aldrich, #H3149), 20 ng/ml EGF (Peprotech, #AF-100-15), 20 ng/ml bFGF (Peprotech, #AF-100-18B) and 100 U/ml penicillin-streptomycin (Fisher Scientific, #11548876).

### Human HepaSH hepatocytes cultures

Human HepaSH hepatocytes were prepared at Central Institute for Experimental Animals (Kawasaki, Japan), as previously described^75^. HepaSH cells derived from one human donor were used in this study. The vial content was transferred into a 50 ml tube containing 40 ml PBS and centrifuged at 200 x *g* for 2 min at 4°C. Cell pellet was then resuspended in recovery medium (Biopredic International, #MIL130) and centrifuged at 200 x *g* for 2 min at 4℃. The cell pellet was then resuspended in seeding medium (Biopredic International, #MIL221) composed of: Williams E GlutaMAX™ supplemented with 100 UI/mL penicillin, 100 µg/mL streptomycin, 4 µg/mL bovine insulin and 10 % fetal calf serum. HepaSH cells were then plated on collagen I-coated plates. After 24 h, medium was discarded, and HepaSH cells were maintained in culture medium (Biopredic International, #MIL222) composed of: Williams E GlutaMAX™, supplemented with 100 UI/mL penicillin, 100 µg/mL streptomycin, 4 µg/mL bovine insulin and 50 µM hydrocortisone. Media were renewed every 2 days. HepaSH cells, media and supplements were kindly provided by Biopredic International.

### Primary mouse cortical neuron cell cultures

Mouse cortical neurons were prepared from E17.5 mouse embryos. Briefly, brains were isolated from embryos under a dissecting microscope, the midbrain and thalamic tissues were gently removed to leave an intact hemisphere containing the cortex and hippocampus. The cortex was then dissected and dissociated using a papain solution (0.5mM of EDTA, 1mM L-Cysteine and 0.4mg/mL of papain (7 units of papain/mL) (Sigma, #76216)) at 37°C in a water bath for 25 min. Neurons purification was performed using the Neuron Isolation kit (Miltenyi, #130-115- 389) according to the manufacturer’s instructions. After purification, neurons were plated at a density of 0.8x10^6^ cells per well of 6-well plates coated with 50 ug/mL Poly-D-Lysine (Thermofisher, #A38904-01) and 1ug/mL Laminin (Sigma, #L2020) in 2mL of Neurobasal media (Fisher Scientific, #11570556) supplemented with 1X B-27 supplement (ThermoFischer, #12587010), 100 U/ml penicillin-streptomycin (Fisher Scientific, #11548876), 2 mM GlutaMAX (ThermoFischer, #35050061). Half of the media was replaced by a fresh media every 2-3 days.

### Ectopic expression of NDI1

The Lenti6.3/V5 NDI1 and pLenti6.3/V5 plasmids were kindly provided by Dr. Joseph Marszalek^59^. HEK293T cells were transfected with 75 µg lentiviral plasmid (pLenti6.3/V5 NDI1 or pLenti6.3/V5), 50 µg PAX2 packaging plasmid and 20 µg of SD11 viral envelope plasmid to produce the viruses. 1 × 10^5^ BTSCs were transduced in a 6-well plate with the virus. After overnight incubation, the virus was washed off and BTSCs were re-plated in BTSC media. Transduced cells were selected through growth in 2.5 µg/ml blasticidin.

### BTSC treatment with mubritinib

BTSCs were dissociated to single cell suspension using Accutase solution (Fisher Scientific, #15323609), plated in low-attachment plates or flasks, and treated with mubritinib (Medchemexpress, #HY-13501) at the time of plating at the indicated concentration, or with vehicle (DMSO 0.1%) for the indicated time points.

### Resipher real-time bioenergetic analysis

To measure oxygen consumption rate (OCR) as an indication of mitochondrial metabolic activity, a Resipher real-time cell analyzer (Lucid Scientific) was used. BTSCs were chemically dissociated using Accutase prior to plating. Cells were plated at a concentration of 1 × 10^5^ cells/well in Nunc 96-well plates (Thermo Fischer Scientific, #10212811) in BTSC media. The plates were then incubated at 37 °C in a humidified atmosphere of 5% CO2 incubator for 1 h. Following incubation, BTSCs were placed on the Resipher system for real-time monitoring of the OCR in absence or presence of 3.0 µM carbonyl cyanide-4- (trifluoromethoxy) phenylhydrazone (FCCP) to measure the basal and the maximal respiration, respectively. For mubritinib-treated conditions, following plating and the 1 h-incubation, mubritinib was added in absence or presence of 3.0 µM FCCP, and subjected to OCR monitoring.

### High-resolution respirometry

Cellular oxygen uptake was quantified by high-resolution respirometry using the Oroboros® Oxygraph-2K (Oroboros Instruments, Innsbruck, Austria). Briefly, 24 h following treatment with mubritinib or vehicle control, BTSCs were chemically dissociated using Accutase, counted and suspended in BTSC medium at a concentration of 2 x 10^6^ cells/ml. 500 ul of the cell suspension were transferred into the Oroboros chamber to allow measurement of the O2 concentration. The cell suspensions were continuously stirred at 750 rpm. Cellular respiration was quantified in terms of oxygen flux based on the rate of change of the O2 concentration in the chambers in absence or presence of FCCP to calculate the maximal respiration.

### MTT assay

The MTT (3-(4,5-dimethylthiazol-2-yl)-2,5-diphenyl tetrazolium bromide) assay was performed to evaluate cytotoxicity of mubritinib^76^. Briefly, the human hepatocytes and mouse neurons were seeded in 96-well plates and treated with various concentrations of mubritinib, in triplicates, for 4 days. After removing culture medium, 100 µl of serum-free medium containing 0.5 mg/ml MTT (Sigma, #M5655) was added to each well and incubated for 2 h at 37 °C. The water-insoluble formazan was dissolved in 100 µl dimethyl sulfoxide, and absorbance was measured at 570 nm.

### Calcein AM cell viability assay

NPCs were plated in 24 well matrigel coated-plates at a density of 4 x 10^4^ cells/well and treated with mubritinib. 4 days after treatment, the number of live cells was estimated using Calcein-AM (ThermoFisher Scientific, #C1430) / DAPI double staining. The cells were incubated at 37 °C for 30 min with 0.5 µM of Calcein-AM and 0.1 ug/mL of DAPI. Images were acquired using a 4X objective on an Eclipe Ti Nikon microscope coupled with NIS analysis software, and a Hamamatsu Digital CCD C10600-10B camera. A minimum of 4 random fields were imaged and counted for each biological replicate. The number of live and dead cells was quantified with Fiji.

### Live cell counting

BTSCs were dissociated to single cell suspension using Accutase. Cells were seeded at a density of 5 x10^4^ cells/well, in low-attachment 6-well plates and treated with mubritinib. Cell number was evaluated 4 days post-plating by the PI cell viability flow cytometry assay using an Accuri C6 flow cytometer.

### Cell death

Cell death was determined using TACS annexin V-FITC apoptosis detection kit (R&D systems, #4830-01-K) according to the manufacturer’s protocol. Briefly, 5 x10^4^ cells/well, were seeded in low-attachment 6-well plates and treated with mubritinib or vehicle control. After 4 days of treatment, co-staining with TACS annexin V-FITC and PI was performed on single cell suspension following the manufacturer’s instructions. Single cell suspension was resuspended in 50 µL of the Annexin V incubation reagent containing 0.5 µL of Annexin V and 5 µL of PI. The fluorescence was analyzed by flow cytometry (Accuri C6 flow cytometer). Data were analyzed using the FlowJo software. Forward and side scatter density plots were used to exclude debris. Doublets were excluded by gating the events in forward scatter height versus forward scatter area. Density plot showing annexin V (X-axis) and PI (Y-axis) staining were used. The quadrant gating was adjusted according to the negative control (unstained cells). Apoptotic cells positive for Annexin V can be seen in the lower right quadrant and dead cells positive for both Annexin and PI in the upper right quadrant. Healthy cells negative for both stains are visualized in the lower left quadrant.

### Ionizing radiation

For measurement of OCR following IR, BTSCs were dissociated to single cell suspension using Accutase. BTSCs were plated in low-attachment cell culture plates or flasks, and then irradiated with 2 or 4 Gy using the Microbeam Technology Irradiation System (XenX Xstrahl®). OCR was assessed 16 h post-irradiation using Resipher Real-time Cell Analyzer (Lucid Scientific).

### Immunoblotting and antibodies

Total proteins were harvested in RIPA lysis buffer containing protease and phosphatase inhibitors (Thermo Fisher Scientific, #A32959). Protein concentration was determined by Bradford assay (Sigma, #B6916), after which samples were subjected to SDS-PAGE and electroblotted onto nitrocellulose membranes. Membranes were blocked in 5% bovine serum albumin in TBST, before sequential probing with primary antibodies and IR-Dye 680 (LI-COR Biosciences, #926-68020) or IR-Dye 800 (LI-COR Biosciences, #926-32212) labeled secondary antibodies. Target proteins were visualized using an Odyssey infra-red scanner (LI-COR). The densitometry of proteins was quantified using Image Studio Lite Software and normalized to tubulin or vinculin. The following antibodies were used: Nestin (1:100, Santa Cruz, #sc-23927), Notch1 (1:1000, Cell Signaling, #CST3608), Olig2 (1:2000, Abcam, #ab109186), SOX2 (1:200, Santa Cruz, #sc-365823), α-tubulin (1:10000, Sigma, #T5168) and Vinculin (1:5000, Invitrogen, #MA511690).

### Extreme limiting dilution assay

Decreasing numbers of BTSCs per well (dose: 25, 12, 6, 3 and 1) were plated in a low attachment 96-well plate, with a minimum of 12 wells/dose, and treated with mubritinib or vehicle control. Seven days after plating, the presence of spheres in each well was recorded and analysis was performed using software available at http://bioinf.wehi.edu.au/software/elda/^77^ to define the percentage of stem cell frequency.

### EdU proliferation assay

BTSCs were dissociated into single cell suspension using Accutase, counted and plated at a density of 5 x 10^4^ cells per well in low-attachment 6-well plate. At the time of plating, BTSCs were treated with mubritinib or vehicle control for 4 days. Then, EdU was added to the culture for 1 h. BTSCs were dissociated into single cell suspension and 3 x 10^4^ cells were plated on 48- well plates coated with poly-D-Lysine (Thermofisher, #A38904-01). When cells adhered to the plate, they were washed with D-PBS, fixed, permeabilized and stained using the Click-iT EdU cell proliferation kit (Thermo Fisher Scientific, #C10340) according to the manufacturer’s protocol. Images were acquired using a 10X objective on an Eclipe Ti Nikon microscope coupled with NIS analysis software, and a Hamamatsu Digital CCD C10600-10B camera. The proportion of cells that incorporated EdU was determined as the ratio of EdU positive cells to the total number of cells.

### Cell cycle analysis

BTSCs were dissociated into single cell suspension using Accutase and 3 x 10^5^ cells were plated in T25 flasks and treated with mubritinib. After 24 h, cells were dissociated to single cell suspension, harvested and fixed with 70% of ethanol overnight at 4°C. The cells were washed with 1X PBS and stained with FxCycle PI/RNase staining solution (Thermo Fisher Scientific, #F10797)^78^. The fluorescence was analyzed by flow cytometry (Accuri C6 flow cytometer). The fractions of G0/G1, S and G2 phase cells were determined using the Watson pragmatic algorithm of FlowJo software.

### Whole-transcriptome analyses (RNA-seq)

BTSC147 was treated with 500 nM of mubritinib for 24 h. Total RNA was extracted with the Qiagen RNeasy Mini Kit according to the manufacturer’s protocol. A DNase treatment of RNA has been added before RNA Cleanup using the RNase-Free DNase set from Qiagen following manufacturer’s instructions. Total RNA quality has been checked using a Fragment Analyzer Systems (Agilent Technologies), RQN were between 8.3 and 10. 100ng of total RNA were converted into Stranded mRNA Library for mRNA-sequencing by using the Illumina Stranded mRNA Ligation kit according to Illumina’s instructions. Briefly, poly(A) tailed RNA are captured by Oligo (dT) beads. The purified poly(A) RNA are fragmented and reverse transcribed into strand complementary cDNA. Then 3’end are adenylated for preparing fragment for the dual indexing adapter ligation. Purified products are selectively amplified for library generation. All libraries have been quantified using a Qubit HS dsDNA kit (Thermofisher) and the library quality check has been performed using a High Sensitivity NGS Fragment Analysis Kit on a Fragment Analyzer System (Agilent Technologies). Indexed libraries were normalized and pooled to be sequenced on an Illumina Novaseq 6000 sequencer according to manufacturer’s instructions with targeted conditions of 2 x 75bp and 25M reads/sample.

### Quantification of mubritinib in the plasma and brain of mice

Following intraperitoneal injection of mice with mubritinib, brains and plasma were collected at different time points. 100 µL of plasma or 100 mg of brain tissue were ground with 100 µL of distilled water using a homogenizer. Then, 600 µL of acetonitrile with [^13^C, ^2^H3]-sorafenib (Darmstadt, Germany) were added and the samples were minced again. Samples were then centrifuged at 10 000 rpm for 5 min. Supernatants were collected and subjected to liquid chromatography / tandem mass spectrometry analysis. Liquid chromatography was performed on an Acquity UPLC system (Waters, USA) linked to the MassLynx software. An Acquity Cortecs C18+ 1.6 µm 2.1 × 50 mm column (Waters, USA) was used. The mobile Phase A was distilled water with acetic acid 1% and formic acid 0.1%; the mobile phase B was acetonitrile. The following gradient was applied: starting with 80% A/ 20% B mobile phases. B was increased to 40% from 0.2 to 0.4 min, to 60% from 0.4 to 0.6 min, to 80% from 0.6 to 0.8 min and to 90% from 0.8 to 1 min, maintained at 90% during 0.75 min, and then reduced to 20% from 2.1 min until the end of the analysis. The flow rate was 0.3 ml/min.

An Acquity TQD detector (Waters, Milford, USA) operated with an electrospray ionization (ESI) source in positive ion mode, using nitrogen as the nebulization and desolvation gas. The ESI ion source was operated at 150°C, with the desolvation temperature at 440°C, cone gas flow adjusted at 50 L/h, desolvation flow at 1000 L/h, and capillary voltage set up at 3.0 kV. MS collision was carried out by argon at 3 × 10^-3^ mBar. Mass spectrometer settings are shown in **Supplementary Table 2**.

### Stereotaxic injections and bioluminescent imaging

For intracranial injections, 3 x 10^5^ patient-derived BTSCs were stereotactically implanted into the right striata (2 mm lateral to the bregma, 1 mm ventral and 3 mm from the pial surface) of 8-week-old male RAGγ2C^−/−^ mice. For the syngeneic model, 5 x 10^5^ luciferase-expressing murine mGB2 cells were into the right striata (2 mm lateral to the bregma, 1 mm ventral and 3 mm from the pial surface) of 8-week-old male C57BL/6N. Mice were randomly assigned to the treatment or vehicle. Seven days post BTSCs injection, mice were administered intraperitoneally with 6 mg/kg mubritinib in 5% DMSO and corn oil suspension or vehicle control 3 days a week (Monday, Wednesday and Friday) until mice were sacrificed. Twelve- and sixteen-days following cell implantation of BTSC73, mice received locally 2 Gy of IR using the Microbeam Technology Irradiation System (XenX Xstrahl®). To examine tumour volume, the animals were intraperitoneally injected with 200 µL of 15 mg/mL D-luciferin (Euromedex, #12507-AATD), anesthetized with isoflurane inhalation, and subjected to bioluminescence imaging using the PhotonIMAGER in vivo imaging system (Biospace Lab). All bioluminescent data were collected and analyzed using M3 vision software. For KM survival plots, mice were collected when they showed signs of tumour related illness.

### Treadmill fatigue test

After 3 months of repeated mubritinib treatment, mice were familiarized and trained to the treadmill apparatus by 2 training sessions on 2 separate days as described^79^. On the first day of training, mice allowed first to freely explore the treadmill for 3 min, then the treadmill was turned on at a speed of 3.0 m/min to allow the mice to begin walking. After ensuring that all mice began walking, a 10 min training session was started using the following parameters: start speed was set at 8 m/min; after 5 min, the speed was increased to 9 m/min; after 7 min, the speed was increased to 10 m/min and the session was stopped at 10 min. On the second day of training, a 15 min training session was performed using the following parameters: start speed was set at 10 m/min; after 5 min, the speed was increased to 11 m/min; after 10 min, the speed was increased to 12 m/min, and the treadmill was stopped after 15 min. For the test, the electric grid shockers that serve to encourage running of the mice were turned on and the test was performed as previously described^79^. At exhaustion, the time and speed were recorded and the distance traversed was calculated. Exhaustion was defined as the point at which mice stopped running and remained on the electric grid shockers for more than 10 s. During the training sessions and the test, the treadmill angle of inclination was set to 5°.

### Immunohistochemical staining and analyses

Immunohistochemical staining of brain tissue sections were performed following paraformaldehyde (PFA) fixation, paraffin embedding, sectioning and staining. Briefly, mice were subjected to transcardiac perfusion with 4% PFA diluted in PBS, then brains were collected and incubated for 24 h in 4% PFA. Brains were then, washed, dehydrated and embedded in paraffin. 5 µm thick sections were performed using a microtome. Sections were then deparaffinized and subjected to antigen retrieval with tris-EDTA buffer (pH 9). Slides were then permeabilized with 0.4% Triton-X100 and blocked with 5% normal donkey serum in PBS-Tween (0.1%). The slides were, then, incubated with the following primary antibody diluted in PBS-Tween (0.1%) for 1 h at RT: anti-Olig2 (1:100, Abcam, #ab109186), anti-Cleaved caspase 3 (1:100, Cell signaling, #D175), anti-SOX2 (1:100, Proteintech, #66411-1). Slides were then washed and incubated with fluorescence-conjugated secondary antibodies (Invitrogen) at RT for 1 h, counterstained with 5 µg/ml of DAPI, mounted using ProLong Diamond Antifade Mountant (Fisher Scientific, #15205739), and imaged using a slide scanner (Hamamatsu Nanozoomer 2.0HT). For quantifications in tumours or healthy host brains, the percentage of positive cells was quantified using Fiji, and the average of 3 fields (minimum 18 x 10^3^ nuclei) on 3 coronal sections (n= 3 mice) is shown. To quantify the neural progenitor cells (NPCs) in the subventricular zone (SVZ), all the nuclei in SVZ are quantified.

To estimate the tumour volume, 5 µm coronal sections were collected every 100 µm over a 600 µm span of the anterior-posterior axis of the brain (n ≥ 3 mice). Hematoxylin-eosin staining was done and whole sections of the brain were imaged using the slide scanner (Hamamatsu Nanozoomer 2.0HT). Tumour volume is calculated using the following equation: (L/3*(A1+ √ (A1*A2)+A2). L is the distance between two sections and A the respective tumour area measured on each section. The volumes between each section are added to obtain the tumour volume on a distance of 600 µm.

### Statistical analysis

Statistical analysis was performed using ANOVA and Student’s t-test, with the aid of GraphPad software 7. Two-tailed and unpaired t-tests were used to compare two conditions. One-way ANOVA with Tukey’s or Dunnett’s post hoc analyses were used for analyzing multiple groups. Data are shown as mean with standard error of mean (mean ± SEM). The log-rank test was used for statistical analysis in the Kaplan-Meier survival plot. p-Values of equal or less than 0.05 were considered significant and were marked with an asterisk on the histograms. p-Values of less than 0.05 are denoted by *, p-values of less than 0.01 are denoted by **, and p-values of less than 0.001 are denoted by *** on the histograms.

## Acknowledgements

This work was supported by grants from Fondation de France (N° 261439), Cancéropôle GSO (N° 2023-E5), Association pour la Recherche sur les Tumeurs Cérébrales (N° 283008), Institut National du Cancer PLBIO (N° 227441 and N° 241284) to AS, AB, TD and AB. AS and AB were supported by a fellowship from the Fondation de France. Cloe Tessier is a recipient of the NewMoon PhD fellowship (Bordeaux University) and Region Aquitaine. We thank staff (Marie-Alix Derieppe, Julie Martineau, Marcia Campistron and Marie-Paule Algeo) at the animal core facility “Animalerie Mutualisee de Talence”, for assistance with studies involving mice. We thank the authors of Molina et al.^59^, for NDI1 plasmids. We thank Dr. Peter Angel for providing the mGB2 cell line. We thank Dr. Deborah Tribouillard-Tanvier for assistant with the Treadmill studies. We thank Dr. Arnaud Mourier for assistance and access to the Oroboros apparatus. We are grateful to Biopredic International (Saint Grégoire, France) and the Central Institute for Experimental Animals (Kawasaki, Japan), for kindly providing the HepaSH cells. We thank Biopredic International for providing all the required media and supplements for HepaSH cells.

---

Supplementary Figure 1. Mubritinib decreases proliferation of patient- and murine-derived BTSCs without induction of cell death. (a) The murine glioblastoma cells, mGB2, were exposed to increasing concentrations of mubritinib (0 to 500 nM) for 4 days, following by live cell counting. (b) The area under the mubritinib dose-response curves (AUC) of wild-type (wt) TP53 BTSCs (#50 and P3) and mutated (mut) TP53 BTSCs (#12, #25, #53, #73 and #147) were plotted. (c) The area under the mubritinib dose-response curves (AUC) of unmethylated MGMT BTSCs (#12, P3 and #147) and methylated MGMT BTSCs (#50, #53, and #73) were plotted. The BTSC25 are not included in this comparison since they are hemi-methylated. (d) The percentages of dead cells (PI positive) and early apoptotic cells (Annexin V positive and PI negative) were measured by Annexin V/PI double staining followed by flow cytometry in BTSCs treated with increasing concentrations of mubritinib for 4 days. One-way ANOVA followed by Dunnett’s test vs vehicle CTL (a), unpaired two-tailed t test (b, c). ***p < 0.001.

Supplementary Figure 2. Gene expression profiling of mubritinib-treated BTSCs. (a-c) Control and mubritinib-treated BTSC147 were subjected to RNA-sequencing analysis. Principal component analysis (PCA) of RNA-seq data is shown (a). Heatmap illustration of hierarchical clustering of the overall gene expression profiles is presented (b). Gene set enrichment analysis of deregulated genes in BTSC147 treated with 500 nM of mubritinib for 24 h demonstrates enrichment for gene sets corresponding to alanine, aspartate and glutamate metabolism-related pathways (c).

Supplementary Figure 3. Mubritinib impairs BTSC stemness properties without affecting the non-oncogenic NSCs. (a) mGB2 cells were subjected to extreme limiting dilution analysis to estimate the stem cell frequencies (SFC) following treatment with increasing concentrations of mubritinib (0 to 500 nM) or vehicle control. (b) mGB2 cells were treated for 4 days with 500 nM of mubritinib or vehicle control and subjected to immunoblotting using the antibodies indicated on the blots. Vinculin was used as loading control. Densitometric quantifications of Olig2 and SOX2 protein level normalized to their corresponding loading control are presented. (c) Non-oncogenic normal NSCs were treated for 4 days with 500 nM of mubritinib or vehicle control and subjected to immunoblotting using the antibodies indicated on the blots. Tubulin was used as loading control. Data are presented as the mean ± SEM, n = 3. One-way ANOVA followed by Dunnett’s vs vehicle CTL (a), Unpaired two-tailed t test (b), *p < 0.05, **p < 0.01, ***p < 0.001.

