## Supplementary figures and images for "Exploiting a metabolic vulnerability in brain tumour stem cells using a brain-penetrant drug with safe profile"

### Supplementary figure 1

#### Supplementary Figure 1

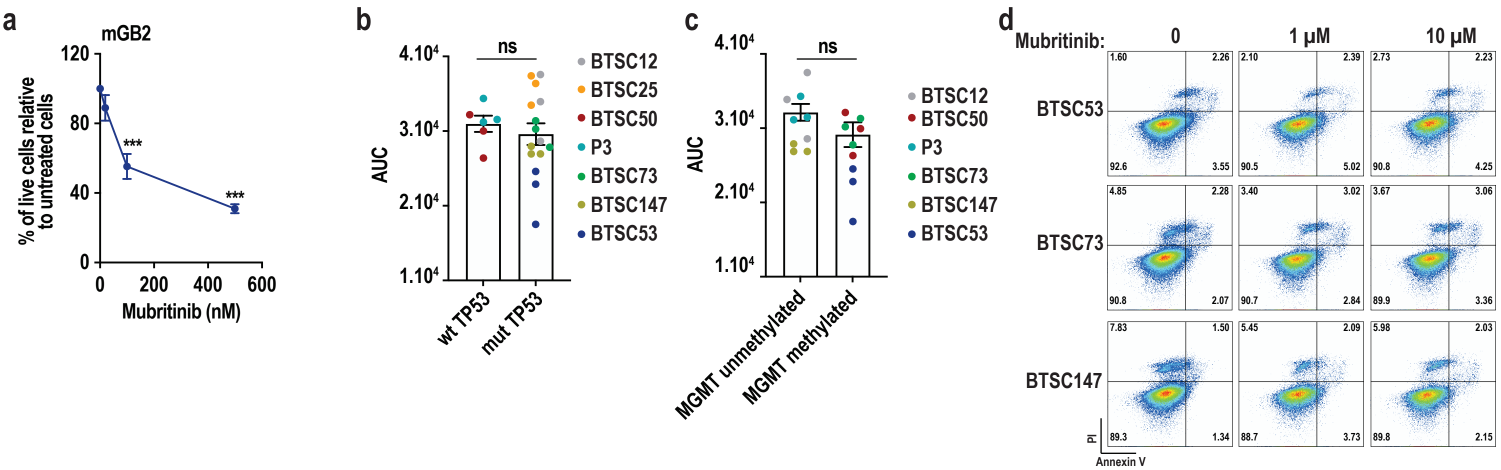

### Supplementary figure 2

**Supplementary Figure 2**

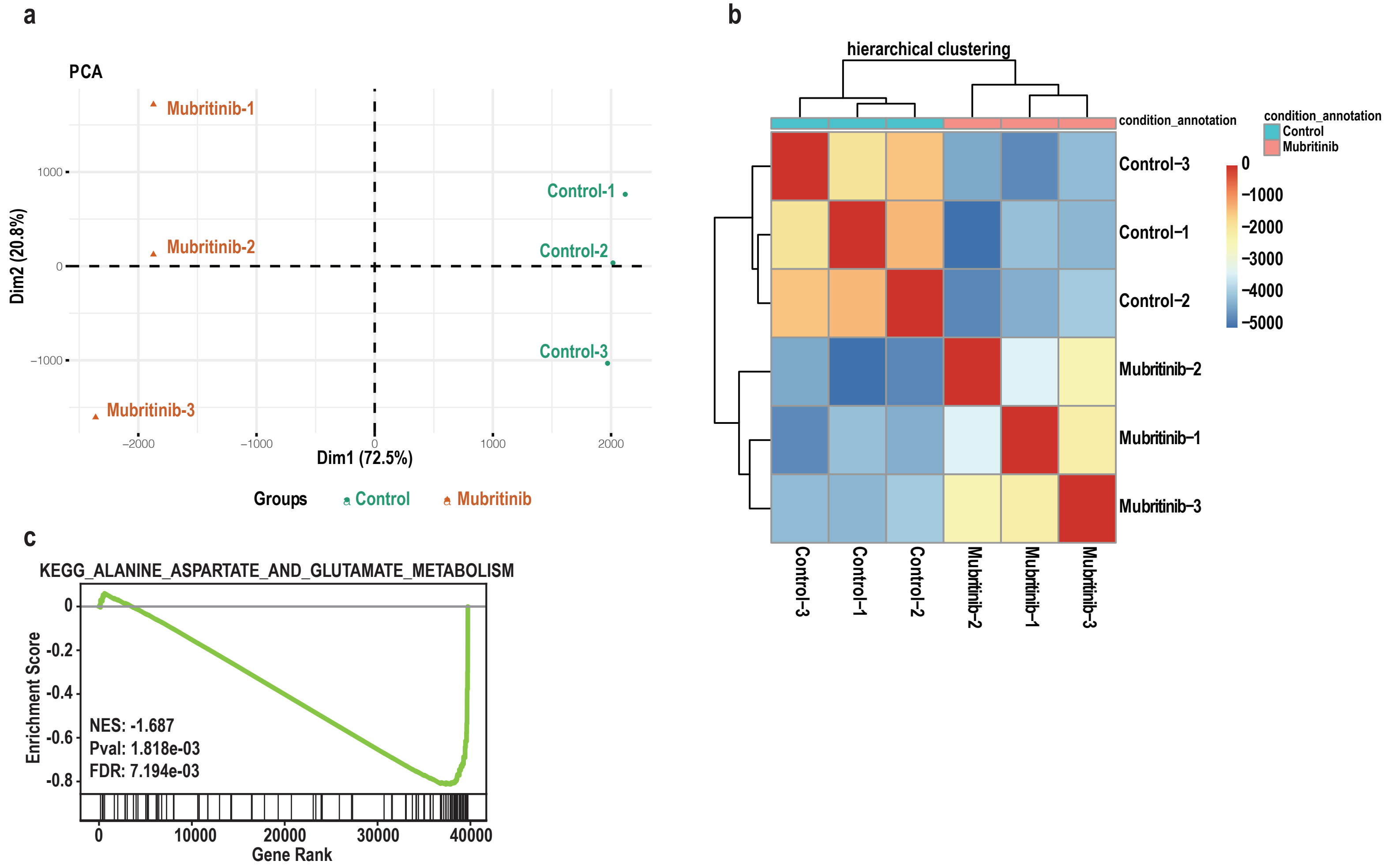

### Supplementary figure 3

#### Supplementary Figure 3

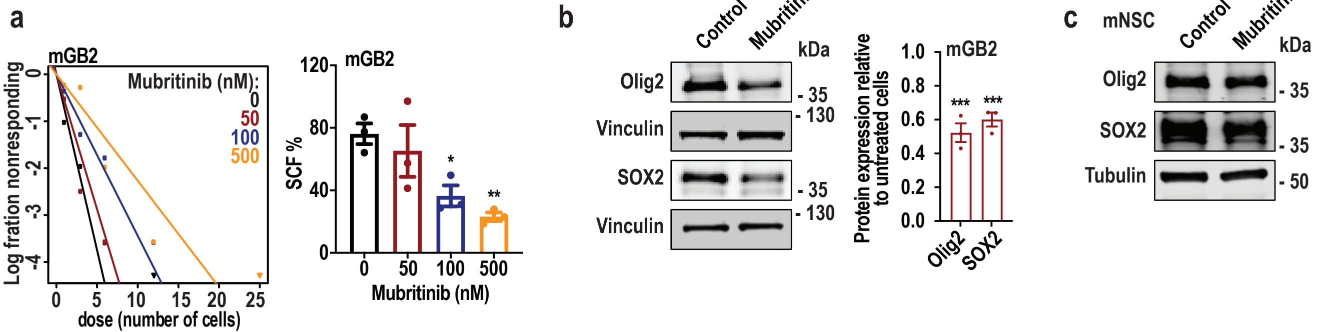
