## Supplementary table 1 for "Exploiting a metabolic vulnerability in brain tumour stem cells using a brain-penetrant drug with safe profile"

**Supplementary Table 1.** Characterization of patient-derived BTSCs

| BTSCs | Diagnosis | EGFR | IDH1 | TP53 | PTEN | NF1 | CDKN2A | MGMT |
|---|---|---|---|---|---|---|---|---|
| **12** | GB-rec[1] | wt[2] | wt[2] | mut[2] | mut[2] | NA[2] | NA[2] | U[1] |
| **25** | GB-rec[3] | wt[2] | wt[2] | mut[2] | mut[2] | NA[2] | NA[2] | U/M[3] |
| **50** | GB[4] | wt[2] | wt[2] | wt[2] | mut[2] | wt[2] | homo del[2] | M[4] |
| **53** | GB[1] | mut[2] | wt[2] | mut[2] | wt[2] | wt[2] | homo del[2] | M[1] |
| **73** | GB[1] | VIII[2] | wt[2] | mut[2] | mut[2] | NA[2] | homo del[2] | M[1] |
| **147** | GB-rec[1] | VIII[2] | wt[2] | mut[2] | mut[2] | wt[2] | homo del[2] | U[1] |
| **P3** | GB[5] | wt[6] | wt[7] | wt[5] | mut[5] | wt[5] | homo del[5] | U |

Mutant (mut) or wild-type (WT) status of genes frequently mutated in GB for the seven BTSC lines used in this study isolated from primary glioblastoma tumour (GB) or recurrent glioblastoma (GB-rec). vIII indicates *EGFR* variant III, homo del indicates a homozygous deletion, NA – not available; U indicates unmethylated; M indicates methylated.
