## Supplementary table 2 for "Exploiting a metabolic vulnerability in brain tumour stem cells using a brain-penetrant drug with safe profile"

**Supplementary Table 2.** Mass spectrometer settings for the quantification of mubritinib by UPLC-MS/MS

| **Analyte** | **Parent ion (m/z)** | **Transition ion (m/z)** | **Confirmation ion (m/z)** | **Cone voltage (V)** | **Collision energy (eV)** |
| --- | --- | --- | --- | --- | --- |
| Mubritinib | 469.2 | 252.3 | 224.2 | 30 | 25 / 20 |
| [^13^C, ^2^H_3_]-Sorafenib | 469.3 | 274.2 | 256.2 | 50 | 30 / 25 |
